# Threonine nitrogen isotopes reveal hidden physiological dimensions of mammalian ecology

**DOI:** 10.64898/2026.08.14.744972

**Authors:** Julia V. Tejada

## Abstract

Stable nitrogen isotopes of amino acids are widely used to reconstruct trophic position. Most applications rely on only two amino acids despite routinely measuring many others. Here, comparative amino acid δ^15^N values from 88 mammal species reveal that threonine records a physiological dimension beyond trophic position. Adding threonine to the canonical glutamate-phenylalanine framework reveals ecological differentiation obscured by broad dietary categories and opposite isotopic relationships between herbivores and secondary consumers. A mechanistic model links this variation to preferential intestinal utilization of threonine for mucin synthesis and predicts experimentally testable patterns of isotope partitioning. These findings show that amino acid δ^15^N values encode complementary dimensions of organismal ecology, expanding amino acid isotope analysis beyond trophic reconstruction to reveal physiological and ecological variation associated with dietary specialization.

## Main Text

The diversification of life has been shaped in large part by how organisms acquire and exploit food resources. Diet structures ecological interactions, governs energy flow through ecosystems, and has repeatedly driven major evolutionary transitions, from the Cambrian emergence of carnivory to the radiation of angiosperms and their herbivores (*1, 2*). Yet because feeding ecology is seldom directly observable in extinct organisms and often difficult to quantify even in living species, reconstructing diet remains a central challenge in ecology and evolutionary biology.

Stable isotopes analysis has transformed this effort by providing biochemical tracers of feeding ecology. Among these approaches, nitrogen isotope analysis of amino acids is particularly powerful because it separates trophic position from environmental variation of nitrogen at the base of the food web (*3*). This ability derives from predictable differences in amino acid metabolism. For example, phenylalanine (Phe) largely retains the isotopic composition of primary producers, whereas glutamic acid/glutamate (Glx) becomes progressively enriched in ^15^N through repeated transamination and deamination reactions (*4*). As a result, the difference in nitrogen isotope ratios (i.e., ‘isotopic offset’ or ΔGlx-Phe) between Glx and Phe has become the canonical proxy for trophic position and has been applied across systems ranging from modern food webs to archaeological assemblages and fossil ecosystems (*5–7*), among other settings.

Despite its success, however, the Glx–Phe framework ultimately reduces feeding ecology to a single dimension defined by these two amino acids (*3, 8*). Yet diet itself is inherently multidimensional, reflecting not only trophic position but also digestive physiology, resource specialization, and metabolism. Routine GC–IRMS analyses typically measure a dozen amino acids simultaneously, raising the possibility that the additional compounds encode independent dimensions of ecological information. This possibility has received little systematic investigation (but see *5, 9*), particularly among terrestrial vertebrates, where comparative datasets remain taxonomically limited and ecological inference continues to rely on linear relationships based on predefined dietary categories.

This study analyzes amino acid δ^15^N values from the largest comparative dataset assembled for continental mammals, more than doubling the existing dataset from 33 to 88 species (175 individuals or pooled samples) across 15 orders and 44 families. Unsupervised learning identified threonine (Thr) as a previously underappreciated physiological axis of isotopic variation that decouples dietary inputs from digestive physiology, improving dietary classification where the canonical Glx–Phe framework fails. More fundamentally, ΔGlx–Thr and ΔGlx–Phe covary in opposite directions in herbivores and secondary consumers, revealing that Thr records a physiological dimension largely independent of trophic position. Together, these findings demonstrate that amino acid δ^15^N values encode multiple, largely independent dimensions of ecology associated with dietary specialization, gut function, and nitrogen metabolism, expanding it from a trophic position proxy to a multidimensional framework for studying animal ecology.

## Multi amino acid δ^15^N reveals ecological structure across mammals

Amino acid δ^15^N measurements were compiled from 88 mammalian species (175 individuals or pooled samples). The dataset comprises 12 amino acids routinely quantified by GC–IRMS analyses (Data SI). The analyses addressed two questions left unresolved by the conventional Glx-Phe approach: whether amino acid δ^15^N can distinguish broad feeding strategies without first classifying specimens by diet, as is typically required when fitting separate linear relationships to groups such as herbivores and secondary consumers, and whether amino acids beyond Glx and Phe encode ecological and physiological information beyond trophic position. To that effect, an unsupervised clustering framework was applied to individual-level amino acid δ^15^N data (species means for published cetacean datasets).

Because the dataset spans taxa from ecosystems with widely varying δ^15^N values at the base of the food web (i.e., that of primary producers), amino acid δ^15^N values were normalized relative to Glx prior to analysis, allowing comparisons to focus on within-sample isotopic differences rather than variation in absolute δ^15^N values among ecosystems. Glutamic acid occupies a central position in nitrogen metabolism, serving as the primary α-amino group donor during transamination reactions (*4, 10*), making it a biochemically grounded reference that reflects the upper bound of nitrogen transfer during amino acid metabolism (*3, 11*). Glx normalization produced better agreement with independently established feeding ecologies than Phe normalization (table S1) and used for results reported in the main text.

Details of the analytical workflow, including script code, missing-data imputation, feature selection, sensitivity analyses, and normalization comparisons, are provided in the Supplementary Information.

## Threonine reveals hidden trophic structure

Comparison across the full amino acid dataset identified Thr as the strongest complement to the canonical Glx–Phe framework. Among the amino acid combinations evaluated, Phe and Thr relative to Glx yielded the highest agreement between unsupervised isotope-based clustering and independently known feeding ecologies (SI). Adding Lys produced similarly high agreement, but nearly half of Lys measurements required imputation and its inclusion did not improve performance relative to the simpler Glx–Phe–Thr feature set. Subsequent analyses therefore focused on Glx, Phe, and Thr.

Incorporating Thr into the canonical Glx–Phe isotope space added a complementary dimension that sharpened trophic resolution (Fig. 1A). Unsupervised clustering of Glx, Phe, and Thr δ^15^N identified ten isotopically distinct groups that organized into three broad trophic assemblages corresponding to herbivores, omnivores, and predators (Fig. 1A, Fig. 2, SI). The expanded isotope space cleanly separated herbivores from secondary consumers, whereas some overlap remained when only Glx and Phe were considered (Fig. 1B). Variable-importance analyses further showed that Thr was a major contributor to dietary classification, particularly by improving discrimination between herbivores and omnivores (Fig. 1B, Fig. 2). This additional resolution emerged from the combined behavior of all three amino acids rather than from any single pairwise isotopic contrast.

**Fig. 1.**
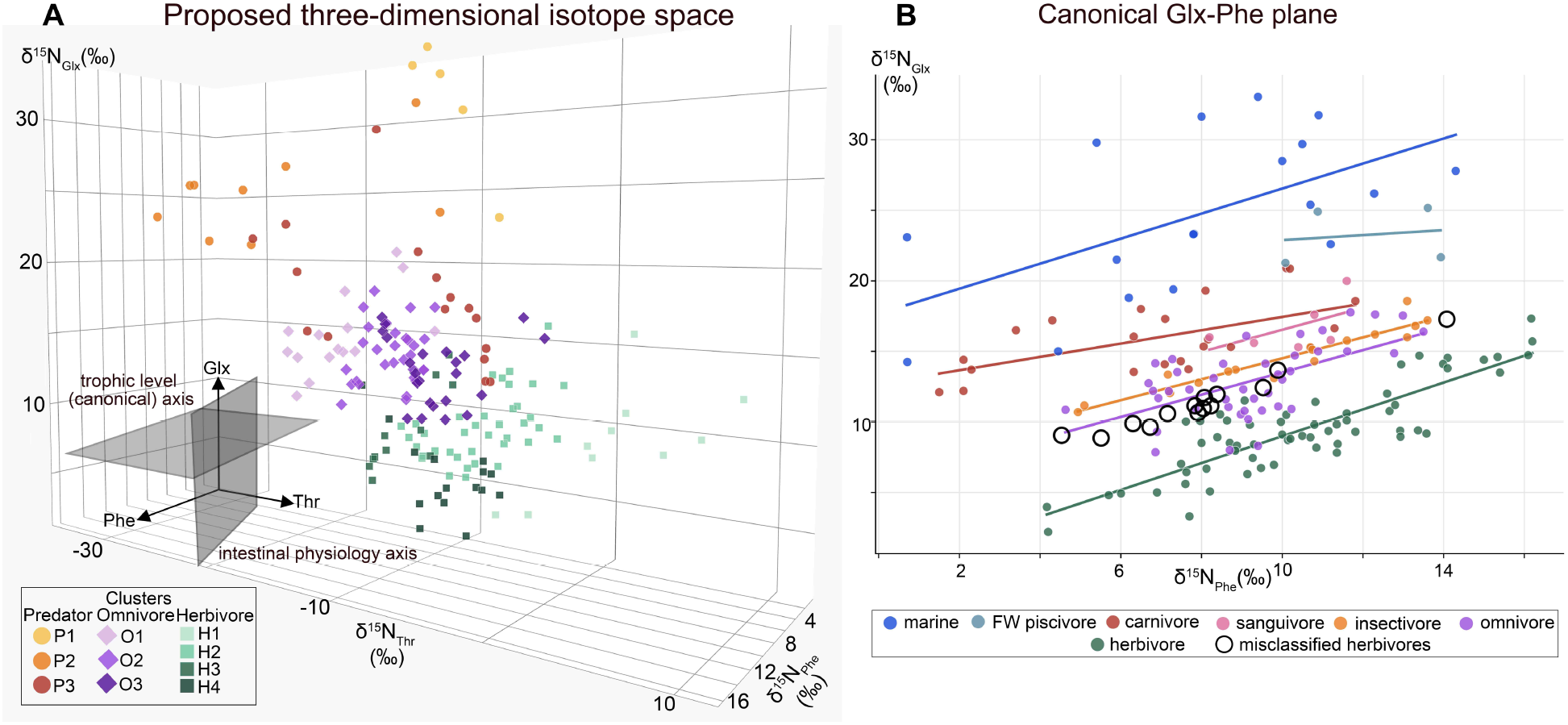
Threonine adds a physiological dimension to canonical amino acid isotope space. (**A**) Three-dimensional isotope space defined by the nitrogen isotope ratios of glutamic acid/glutamine (δ^15^N_Glx_), phenylalanine (δ^15^N_Phe_), and threonine (δ^15^N_Thr_). Symbols denote trophic guilds: circles, predators (P); diamonds, omnivores (O); squares, herbivores (H). (**B**) Within the canonical Glx–Phe δ^15^N framework, linear discriminant analysis misclassified several obligate herbivores (black open circles) as secondary consumers, illustrating the limited discriminatory power of Glx and Phe alone.

**Fig. 2.**
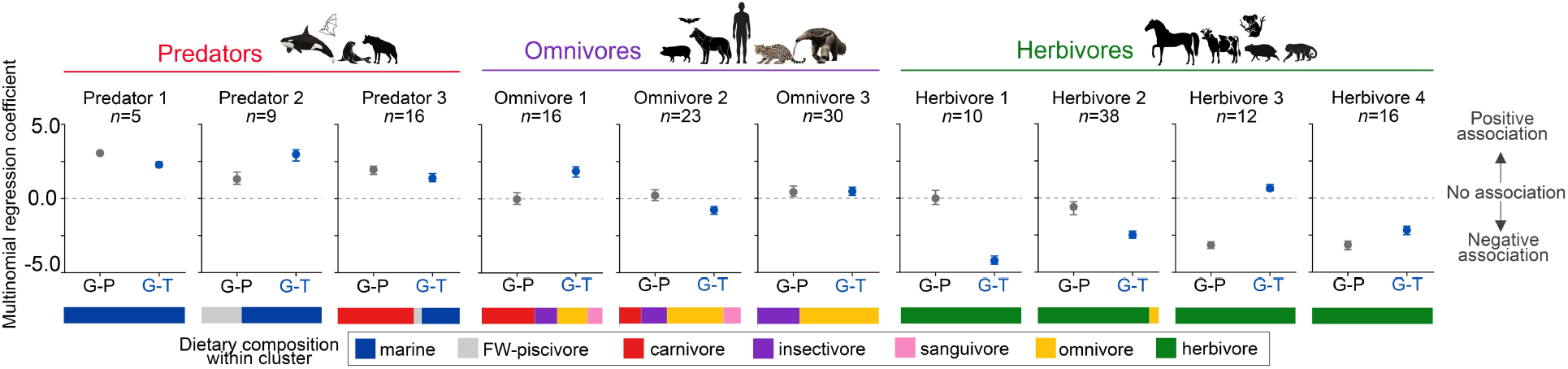
Threonine disproportionately contributes to discrimination among dietary clusters. Multinomial logistic regression fitted to cluster assignments from the unsupervised analysis identified ΔGlx–Thr as a strong discriminator of dietary strategy, particularly among herbivore clusters, where the canonical Glx-Phe framework contributed comparatively little to their separation from omnivores. Points represent the mean standardized multinomial logistic regression coefficients across 20 stratified bootstrap replicates; error bars indicate 95% bootstrap intervals (2.5th–97.5th percentiles). Coefficients near zero indicate little contribution to cluster discrimination, whereas larger absolute coefficients indicate greater influence. Positive and negative values denote the direction of association with each cluster. G–P= glutamic acid– phenylalanine δ^15^N offset, G–T= glutamic acid–threonine δ^15^N offset. Taxonomic composition of each cluster is shown in fig. S1.

The added discriminatory power of Thr was most apparent among herbivores. Under the canonical Glx–Phe framework, fourteen individuals representing six herbivorous taxa exhibited ΔGlx–Phe δ^15^N values exceeding 2.5‰ (table S2), a conservative threshold typically associated with secondary consumers (*12*). These included five rodents (*Hydrochoerus hydrochaeris, Dasyprocta variegata, Sciurus spadiceus, Tamiasciurus hudsonicus*, and *Proechimys* spp.), one sirenian (*Trichechus manatus*), and one pilosan (*Choloepus hoffmanni*). When treated as unknown, linear discriminant analysis trained on Glx–Phe δ^15^N (excluding these individuals from the training dataset) classified all fourteen individuals as omnivores (Fig. 1B, table S2-S3).

Although some of these species may occasionally ingest animal-derived material (*13, 14*), plants overwhelming dominate their diets, making their classification as omnivores ecologically misleading.

The unexpectedly high ΔGlx–Phe δ^15^N values observed in a subset of obligate herbivores are well illustrated by the capybara (*Hydrochoerus hydrochaeris*). Despite feeding almost exclusively on aquatic and semi-aquatic C_4_ grasses (*15*), capybaras exhibit ΔGlx–Phe values (3.4‰–3.7‰) characteristic of omnivores (Fig. 1B). Elevated nitrogen isotope ratios of aquatic settings are unlikely to explain this pattern because they should affect Glx and Phe proportionally and therefore exert little effect on their offset. Instead, cecotrophy, a widespread strategy in capybaras that recycles nitrogen-rich compounds through repeated digestion (*16, 17*), offers a plausible explanation. Repeated transamination and deamination preferentially enrich trophic amino acids such as Glx while leaving Phe largely anchored to dietary baseline values, so sustained coprophagy is predicted to inflate apparent trophic enrichment. Diet-feces δ^15^N offsets approaching 4‰ (*18*), together with documented ^15^N enrichment of cecal nitrogen pools (*19*), suggest that intensive nitrogen recycling can shift obligate herbivores into Glx–Phe space typically occupied by secondary consumers. These observations indicate that ΔGlx–Phe δ^15^N integrates not only primary dietary composition but also the extent of internal nitrogen recycling, complicating trophic interpretation when digestive reprocessing is substantial.

By adding an independent physiological axis, Thr distinguished dietary inputs from digestive processing and revealed substantially greater ecological structure among consumers (Fig. 1A, Fig. 2). Predators and omnivores, for instance, partitioned into three groups, whereas herbivores formed four distinct clusters, revealing previously unrecognized ecological differentiation within primary consumers. Importantly, in a few cases, the unsupervised isotope-based groups did not conform to conventional dietary classifications. Such discrepancies need not represent misclassification: dietary categories summarize predominant feeding habits, but impose discrete boundaries on species that may exploit broader or overlapping dietary niches. For example, one group (Herbivore 2) contained approximately 90% conventionally classified herbivores together with Pallas’s long-tongue bat and peccaries, both broadly classified as omnivores but capable of subsisting largely or entirely on plant resources. The unsupervised framework therefore reveals ecological similarities that can cut across conventional dietary boundaries. Although most conspecific individuals clustered together, some species spanned multiple clusters within the same trophic category. Squirrel monkeys, for example, remained confined to omnivore groups, whereas three-toed sloths partitioned between two herbivore clusters, indicating that amino acid δ^15^N values reveal ecological heterogeneity within trophic guilds while preserving overall food-web structure.

## Threonine captures a physiological dimension beyond trophic information

Despite both being dietary essential amino acids, Phe and Thr encode distinct physiological dimensions of nitrogen metabolism. This distinction is evident in the opposite relationships between ΔGlx–Phe and ΔGlx–Thr in herbivores and secondary consumers (OLS slopes: -0.55 and +2.21; SMA slopes: −1.45 and +2.58 respectively; Fig. 3A). Among secondary consumers, ΔGlx-Thr covaries positively with ΔGlx-Phe (Fig. 3B, SI), consistent with previous observations that Thr δ^15^N decreases with trophic level and increasing dietary protein content (*11, 20, 21*). In herbivores, however, this relationship reverses, with a consistent negative relationship that remains remarkably stable across multiple sensitivity analyses (Fig. 3A, table S4-S5). Moreover, the range of ΔGlx-Thr variation in herbivores (∼16.8‰) is nearly twice that of ΔGlx–Phe (∼8.9‰; measured values only, Fig. 3B), suggesting that Thr captures a substantial source of isotopic variation in herbivores largely independent of the trophic-position signal encoded by Glx–Phe.

**Fig. 3.**
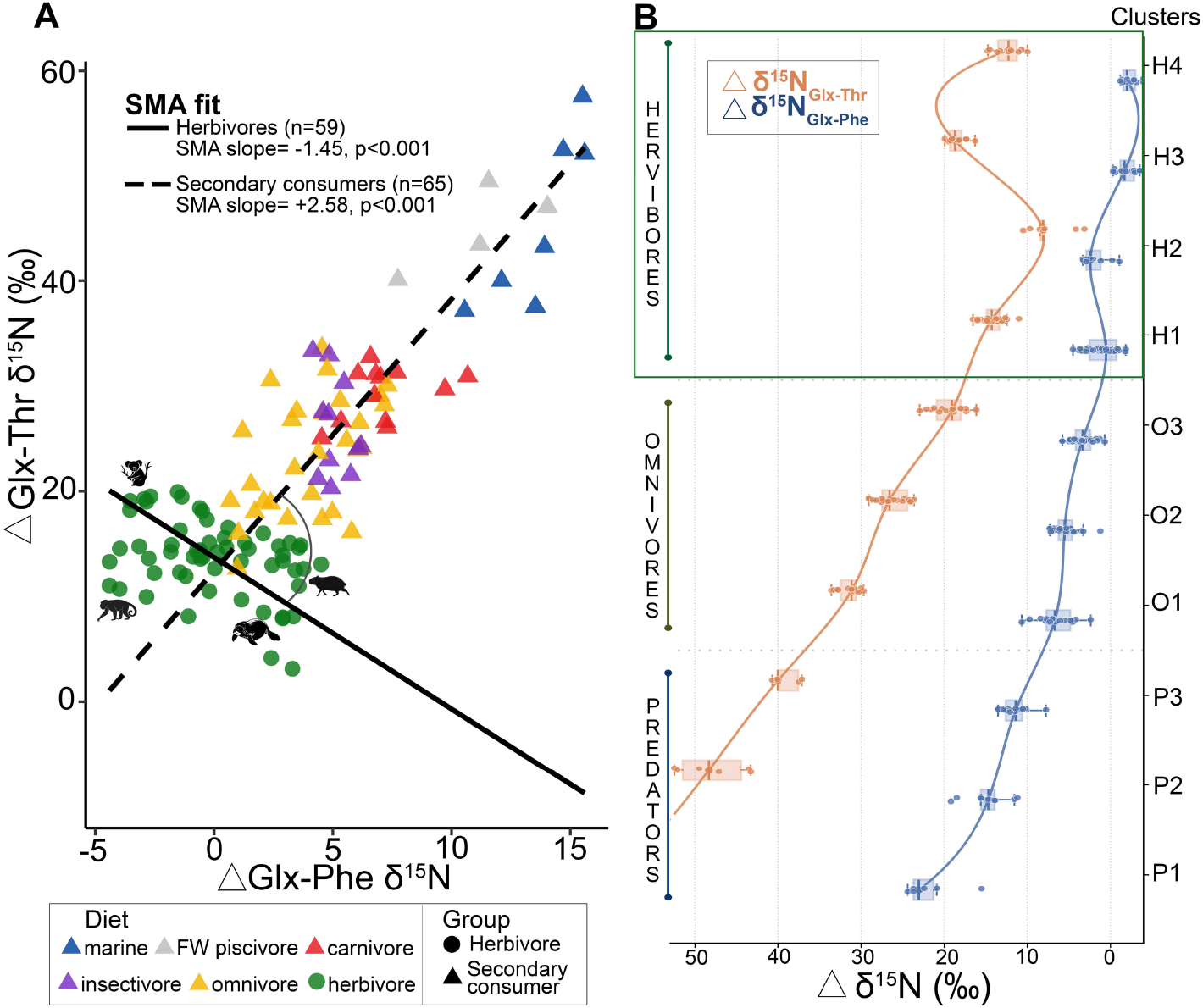
Herbivores and secondary consumers exhibit contrasting isotopic scaling relationships. (**A**) Herbivores and secondary consumers follow opposing scaling relationships in ΔGlx–Phe versus ΔGlx–Thr space, producing nearly orthogonal trajectories. Standardized major axis (SMA) slopes were −1.45 for herbivores and +2.58 for secondary consumers. The SMA slopes differed significantly between the two groups (likelihood ratio test, *P* < 0.001). Silhouettes indicate the positions of four representative herbivore species (koala, capybara, howler monkey, and manatee) along the herbivore continuum. (**B**) Isotopic compositions of dietary clusters reveal contrasting trends: ΔGlx–Thr covaries positively with ΔGlx–Phe across secondary consumers but shows a consistently negative relationship among herbivores. The contrasting relationships remained robust across multiple sensitivity analyses (SI). Curves connect cluster medians and are shown only as visual guides.

Dietary protein availability alone cannot readily explain this variation. If Thr primarily reflected dietary protein content among herbivores, specialized folivores such as koalas would be expected to occupy the opposite end of the observed gradient. Instead, koalas exhibit some of the most ^15^N-depleted Thr values among herbivores, overlapping those of several omnivores (Cluster H3 in Fig. 3B). This pattern points instead toward intrinsic physiological differences associated with herbivory. Because herbivores differ markedly in digestive physiology, fermentation strategy was tested as a potential explanation to their large ΔGlx-Thr variation. Hindgut fermenters exhibited a negative relationship between ΔGlx-Thr and ΔGlx-Phe, whereas the relationship was weaker among foregut fermenters (SI). However, SMA detected no significant difference between slopes (likelihood ratio= 3.16, P= 0.075, table S5), indicating that fermentation strategy alone does not account for the observed patterns.

The contrasting scaling relationships also persisted after accounting for uneven phylogenetic sampling. SMA analyses were repeated across 1000 family-thinned datasets, each generated by randomly retaining one representative per mammalian family within herbivores and secondary consumers independently. Herbivore slopes remained negative and secondary consumer slopes positive in every resampled dataset, with median slope estimates nearly identical to those obtained from the complete dataset (Table S6). These analyses demonstrate that the contrasting isotopic scaling relationships are not driven by overrepresentation of particular mammalian families.

## A mechanistic hypothesis linking intestinal threonine utilization to δ^15^N variation

A parsimonious explanation for the contrasting relationships between ΔGlx-Phe and ΔGlx-Thr among consumers arises from the distinctive metabolism of Thr within the mammalian intestine. Among essential amino acids, Thr is unusual in being extensively utilized during intestinal first-pass metabolism, where as much as 80% of dietary Thr may be retained before reaching systemic circulation (*22, 23*). The exceptional intestinal demand for Thr reflects its central role in biosynthesis of mucins, the heavily glycosylated proteins that protect and lubricate luminal epithelial surfaces (*24*–*27*). Mucin peptide backbones are exceptionally rich in Thr, Ser, and Pro, with Thr comprising up to ∼35% of the polypeptide sequence in some mucins (*28*).

I propose a model whereby disproportionate intestinal demand partitions dietary Thr into two isotopically distinct pools (Fig. 4). Under this model, preferential transport of lighter Thr isotopologues (i.e., ^14^N-Thr) across the intestinal epithelium would enrich the residual intestinal Thr pool in ^15^N, providing the substrate for mucin biosynthesis, while peripheral tissues receive isotopically lighter Thr. Similar kinetic isotope effects have been demonstrated for biological membrane transport systems in mammals (*29, 30*). Although isotope fractionation during intestinal amino acid transport has not been measured directly, several observations are consistent with the predicted tissue partitioning (*31*). Preliminary measurements obtained for this study show that intestinal mucus from a chicken was enriched in ^15^N-Thr relative to feather, a peripheral tissue, by an amount exceeding analytical variability (fig. S2). The intestinal mucin retention hypothesis predicts that Thr in downstream peripheral tissues (e.g., bone, muscle, etc.) should be more ^15^N-depleted than Thr in tissues with higher amino acid turnover, such as blood and liver, a pattern reported in both captive kestrels and wild herring gulls (*31*). Isotopic mass balance further predicts that the magnitude of Thr ^15^N depletion in peripheral tissues should increase with the proportion of dietary Thr retained by the intestine. Additional paired measurements of Thr δ^15^N in intestinal mucins and peripheral tissues across species will be required to test these predictions and establish the generality of the proposed mechanism.

**Fig. 4.**
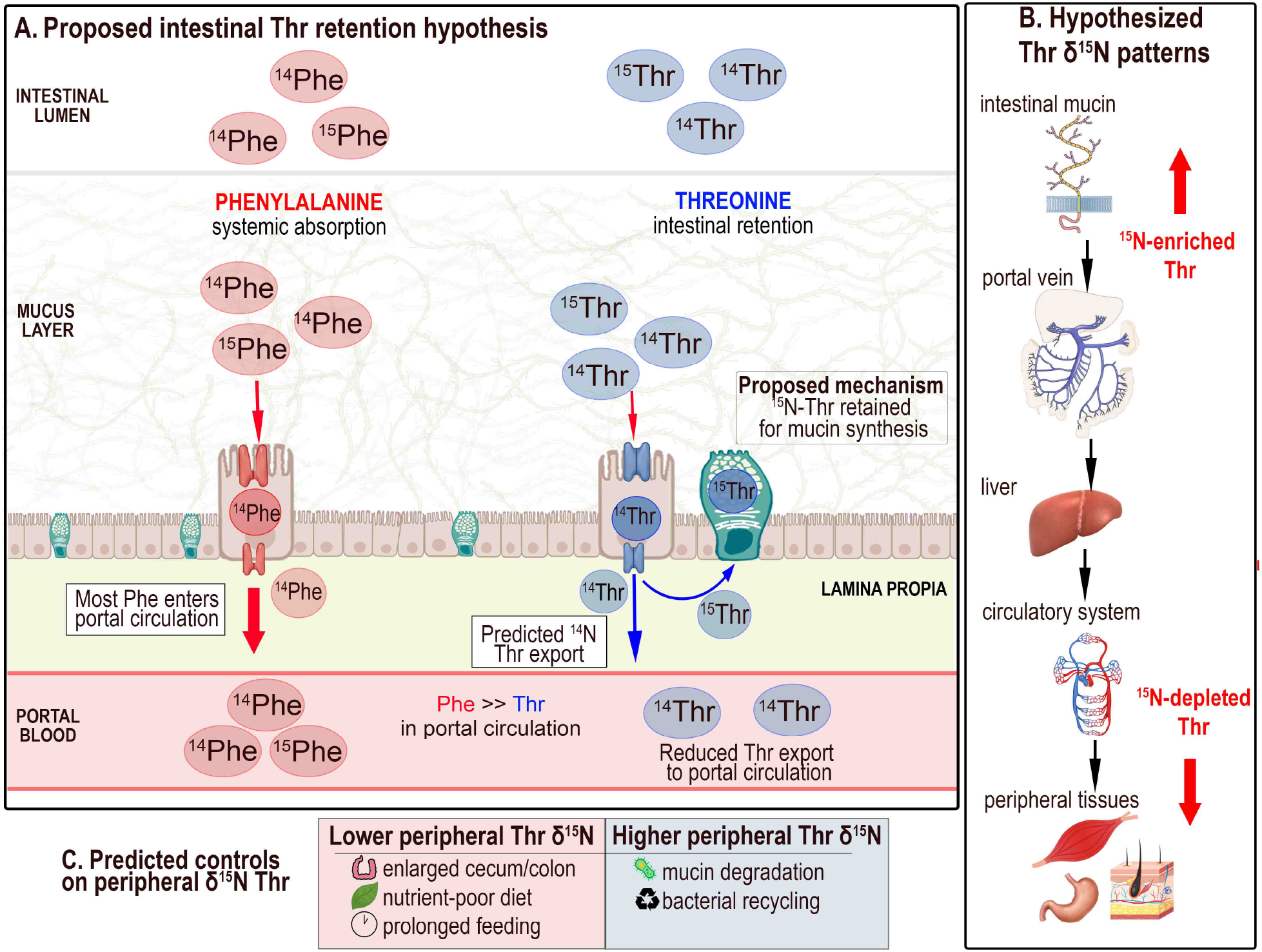
A proposed intestinal Thr retention mechanism for mammalian Thr δ^15^N variation. (**A**) Dietary Thr is hypothesized to partition into two isotopically distinct pools: ^14^N-Thr is preferentially exported to the portal vein for tissue synthesis and catabolism, whereas ^15^N-Thr remains in the intestine and is used by goblet cells for mucin synthesis. (**B**) The model predicts that intestinal mucins become enriched in ^15^N relative to portal blood and peripheral tissues. (**C**) Tissue Thr δ^15^N reflects the balance between two opposing processes: intestinal Thr retention, which drives ^15^N depletion of peripheral tissues, and microbial nitrogen recycling, which partially restores ^15^N-enriched Thr to the systemic amino acid pool. Variation in the relative strength of these processes could generate the broad range of Thr δ^15^N values observed among mammals.

Alternative metabolic mechanisms could also contribute to Thr nitrogen-isotope fractionation, but available evidence appears less consistent with the observed patterns. For example, transamination is unlikely to play a major role in Thr’s δ^15^N because, unlike Glx, Thr does not normally undergo transamination. Catabolism through threonine deaminase (also known as Thr dehydratase) could contribute to diet-tissue isotope fractionation (*32*), but intestinal first-pass utilization greatly exceeds Thr catabolic fluxes (33–*35*). Moreover, *in vitro* experiments using purified Thr indicate that Thr deaminase activity does not generate the observed ^15^N-depletion of catabolic products (*36*).

The intestinal mucin retention hypothesis does not exclude contributions from other processes known to influence Thr δ^15^N, including nutritional state, immune activation, and host-microbiome interactions (*21, 22, 24*). Rather, it provides a framework that may reconcile the trophic and physiological components of the Thr nitrogen isotopic signal. Across consumers, repeated intestinal retention of ^15^N at successive trophic transfers provides a mechanism for the systematic trophic ^15^N depletion of Thr observed in natural and controlled feeding studies (*20, 21, 37*). Superimposed upon this trophic trend, variation in intestinal Thr demand among species could generate the additional isotopic variation observed within trophic groups (*22, 28*). Therefore, unlike Glx, whose isotopic composition primarily reflects nitrogen transfer through transamination, Thr appears to record a physiological dimension of nitrogen allocation associated with intestinal function.

This framework generates several experimentally testable predictions (Fig. 4B-C). Threonine in intestinal mucins should be enriched in ^15^N relative to dietary and circulating Thr, with corresponding ^15^N depletion of peripheral tissues. Experimental manipulation of mucin production should alter tissue Thr δ^15^N among herbivores, and germ-free animals should exhibit altered Thr isotope patterns owing to the absence of microbial nitrogen utilization and recycling (*38*). Together, these predictions provide a direct path toward experimental evaluation of the proposed mechanism.

## Gut morphology and nitrogen recycling modulate threonine δ^**15**^N among herbivores

The exceptionally large range of ΔGlx-Thr observed among herbivores (Fig. 3) is proposed to reflect the remarkable diversity of herbivore digestive anatomies and nitrogen recycling strategies. Herbivores differ substantially in intestinal size, mucosal surface area, gut residence time, and dependence on microbial fermentation (*39, 40*), all of which are expected to influence the partitioning of Thr between intestinal tissues and systemic circulation. Larger intestinal surface areas require greater mucin production to maintain epithelial integrity (*27, 41*). Goblet cell abundance increases progressively along the intestinal tract (*41*), suggesting that species with enlarged ceca or colons should devote proportionally more dietary Thr to mucin biosynthesis. Herbivores also experience greater mechanical abrasion of the intestinal epithelium owing to fibrous diets, while prolonged feeding of nutrient-poor food requires sustained mucus production and epithelial turnover (*41, 42*). Together, these factors are predicted to increase intestinal sequestration of Thr and amplify isotopic ^15^N depletion of peripheral tissues (Fig. 4C).

At the same time, microbial degradation of shed mucins and goblet cells is expected to return ^15^N-rich Thr to the host. Consequently, tissue Thr δ^15^N should reflect the balance between two opposing processes: intestinal retention, which drives ^15^N depletion of peripheral tissues, and microbial nitrogen recycling, which partially restores ^15^N-enriched Thr to the systemic amino acid pool. Variation in either process could therefore generate the broad isotopic diversity observed among herbivores.

Koalas illustrate one extreme of this proposed continuum. Their exceptionally elongated hindgut (*39*), implies unusually high mucin biosynthetic demands, while their specialized *Eucalyptus* diet is associated with markedly reduced gut microbial diversity relative to more generalist herbivores (*43*). Reduced microbial recycling combined with elevated intestinal Thr demand provides a plausible explanation for the very negative Thr δ^15^N values observed in this species (Fig. 2-3). More broadly, herbivores possessing more diverse and metabolically active gut microbial communities are predicted to exhibit more positive Thr δ^15^N values than species with less developed microbiomes, as microbial recycling increasingly offsets intestinal Thr sequestration.

## Implications for amino acid isotope ecology

These results suggest that amino acid δ^15^N values encode complementary dimensions of organismal ecology rather than providing redundant estimates of trophic position. Glx primarily reflects nitrogen transfer through transamination, Phe records environmental nitrogen at the base of the food web, and Thr is hypothesized to record the allocation of dietary nitrogen to intestinal function. Together, these amino acids capture complementary aspects of organismal physiology that become accessible through multi-amino-acid isotope analysis.

Beyond providing a mechanistic explanation for Thr isotope variation, this study shows that expanding the canonical Glx-Phe framework reveals ecological structure obscured when feeding ecology is reduced to discrete dietary categories. The separation of obligate herbivores from omnivores demonstrates this additional resolution. Amino acid isotope ecology can therefore move beyond asking where an organism feeds in a food web toward asking how it acquires and physiologically processes nutrients. If the proposed mucin-retention mechanism is supported by further experimental testing beyond the preliminary evidence presented here, Thr δ^15^N could provide not simply an additional trophic tracer, but a window into intestinal nitrogen allocation and its relationship to dietary specialization.

In herbivores, the broad range of ΔGlx–Thr values and their partitioning into multiple isotopic groups suggest substantial physiological differentiation within a single trophic level. What determines an herbivore’s position along this isotopic continuum remains unresolved, but differences in intestinal morphology, microbial nitrogen recycling, dietary composition, and digestive strategy provide testable possibilities for tracing physiological specializations associated with herbivory. If calibrated in living species, these relationships could allow amino acid isotope patterns to reconstruct not only where consumers sit in a food web, but also the diversity of physiological strategies through which animals exploit resources. Applied across modern and fossil communities, this approach could reveal changes in the diversity of feeding strategies and digestive specialization, providing a more multidimensional view of ecosystem structure than conventional food-web reconstruction.

## Acknowledgments

I am indebted to R. Baptista (U. of Toronto) for advisory and guidance on the statistical framework applied in this study, R. MacPhee (AMNH), V. Pacheco (MHN-UNMSM), J. Nations, V. Mathis (FLMNH), S. Robson, Kayce Bell (NHMLAC) for access to collection specimens under their care, F. Moreno for sample processing assistance, A. Rodriguez for pilot data of intestinal Thr δ^15^N, and J. Eiler for fruitful discussions.

## Data and materials availability

All data are available in the main text or the supplementary materials.

## Supplementary Materials

Materials and Methods

Supplementary Text

Figs. S1 to S2

Tables S1 to S6

Data S1 to S2

