## Supplementary Materials for "Threonine nitrogen isotopes reveal hidden physiological dimensions of mammalian ecology"

Julia V. Tejada

Division of Geological and Planetary Sciences, California Institute of Technology, Pasadena, CA 91125.

**This PDF file includes:**

Materials and Methods
Fig. S1-S2
Tables S1 to S6
Captions for Data S1 to S2

**Other Supplementary Materials for this manuscript include the following:**

Data S1 to S2

### Materials and Methods

#### Samples

Results presented are based on amino acid specific $\delta^{15}N$ data of hair keratin and/or bone collagen from 88 mammal species (175 individuals) spanning 15 orders in 44 families (Data S1). Clustering analyses were performed at the individual level; for marine cetaceans, represented here only by published data, species averages were used. Samples come from the following museum collections: American Museum of Natural History (AMNH), Museo de Historia Natural-UNMSM (MUSM), Natural History Museum of LA County (LACM), and the Florida Museum of Natural History (FLMNH). All applicable international, national, and/or institutional sampling methods were carried out in accordance with relevant guidelines and regulations.

#### Isotopic Analysis

Twelve amino acids were consistently measured across samples: alanine (Ala), glycine (Gly), threonine (Thr), serine (Ser), valine (Val), leucine (Leu), isoleucine (Ile), proline (Pro), aspartic acid/asparagine (Asx), glutamic acid/glutamine (Glx), phenylalanine (Phe), and lysine (Lys). Tyrosine (Tyr), arginine (Arg), and hydroxyproline (Hyp) were were measured in only a subset of samples and were therefore excluded from further analyses. "Big delta" values ($\Delta$) denote isotopic offsets between two amino acids; for example, $\Delta$Glx–Phe, represents the difference between the $\delta^{15}N$ values of Glx and Phe.

Amino acids were chemically isolated and purified from sample material prior to derivatization in preparation for GC-IRMS analysis, following procedures described in (*1*). Samples were analyzed using a Thermo Scientific Delta Q isotope ratio mass spectrometer coupled to a Thermo Trace 1310 GC gas chromatograph through a Thermo GC Isolink II interface. Two internal reference compounds, L-2-Aminoadipic acid (AAA) and L-(+)-Norleucine (Nor), with known nitrogen isotopic composition, were co-injected with each sample and used to assess accuracy and precision. Each sample was analyzed in triplicate. Between triplicate sample analyses, a suite of 14 amino acids with known isotopic compositions were analyzed: Ala, Gly, Thr, Ser, Val, Leu, Ile, Pro, Asx, Glx, Phe, Tyr, Lys, and Arg. AAA and Nor were also co-injected with these amino acid standards. Isotopic correction of unknown amino acids was based on linear calibration relationships derived from the amino acid standard suites analyzed immediately before and after each set of triplicate sample measurements. These corrections were then applied to the measured isotope ratios. All amino acids are reported in $\delta$ notation relative to atmospheric $N_2$. Analytical precision for each sample, expressed as the standard deviation (SD) of triplicate measurements, is reported in Data S1.

#### Unsupervised agglomerative clustering

Samples were grouped using hierarchical agglomerative clustering implemented in scikit-learn (Python). Clustering was based on Euclidean distances and Ward linkage, which iteratively merges observations to minimize within-cluster variance. The number of clusters was not specified *a priori*. For each analysis, an initial clustering model was used to generate the complete agglomeration tree, from which dendrograms and the distribution of Ward linkage distances were obtained. Distance thresholds were then selected by visual inspection of the dendrograms and linkage-distance profiles to identify informative and stable partitions while avoiding excessive subdivision, and clustering was repeated using the selected threshold. Final distance thresholds were 18 for the $\Delta$Glx–Phe plus $\Delta$Glx–Thr analysis (Fig. 1), 8 for analysis using ΔGlx–Phe, and 8 for ΔGlx–Thr alone. The Python script used for all clustering analyses is provided in Data S2.

Missing $\delta^{15}N$ values were imputed prior to clustering using multivariate iterative imputation (IterativeImputer, scikit-learn). For the reduced feature sets, imputation was performed with a maximum of 10 iterations and a fixed random seed (random_state = 0); the complete amino acid dataset was imputed using a maximum of 100 iterations. Missing values varied among amino acids (Ala 12.6%, Gly 6.3%, Thr 28.6%, Ser 26.9%, Val 6.9%, Leu 20.0%, Ile 25.7%, Pro 9.7%, Asx 33.7%, Glx 0%, Phe 0%, Lys 46.3%).

To assess whether the imputation of Thr influenced the clustering results, analyses were repeated using only specimens with measured Thr values, yielding a reduced dataset of 127 observations (Data S1). Within this dataset, missingness for the remaining isotopic offsets was: Glx–Ala 0.0%, Glx–Gly 0.0%, Glx–Thr 0.0%, Glx–Ser 8.7%, Glx–Val 0.0%, Glx–Leu 8.7%, Glx–Ile 11.8%, Glx–Pro 0.8%, Glx–Asx 8.7%, Glx–Phe 0.0%. Repeating the clustering analyses on this dataset produced highly similar cluster assignments (Table. S4), indicating that the principal clustering patterns (Fig. S1) were not driven by imputation of missing Thr values.

The relative importance of isotopic variables for reproducing cluster assignments was also evaluated using random forest classifiers. For each analysis, specimens were randomly divided into training (80%) and test (20%) datasets, and a random forest containing 100 trees was fitted to the training data. Classification accuracy was evaluated on the held-out test dataset and feature importance scores were recorded for each predictor. This procedure was repeated across 20 independent train/test splits.

Throughout the manuscript, amino acid $\delta^{15}N$ values are expressed relative to glutamate/glutamic acid (ΔGlx–AA). Analyses using phenylalanine as the reference amino acid produced qualitatively similar results (Table S1), although Glx normalization provided improved separation among herbivore clusters. Including lysine yielded comparable clustering results but was not adopted for the primary analyses because nearly half (46.3%) of lysine measurements required imputation.

<u>Feature contributions to cluster discrimination were evaluated using multinomial logistic regression</u>

Cluster assignments generated by the hierarchical agglomerative clustering were used as response classes, and isotopic offsets (ΔGlx–Phe and ΔGlx–Thr) were used as predictor variables. To assess the stability of feature coefficients, 20 stratified bootstrap datasets were generated by resampling specimens with replacement within each cluster while preserving the original cluster sample sizes. Predictor variables were standardized within each bootstrap replicate prior to model fitting. A multinomial logistic regression model was fitted using the LBFGS solver. Standardized regression coefficients were recorded for each predictor and cluster. For Fig. 2, points represent the mean coefficient across bootstrap replicates and error bars represent the 2.5$^{th}$ and 97.5$^{th}$ percentiles of the bootstrap distribution. The Python script used for this analysis is provided in Data S2.

<u>Evaluation of herbivore classification using the canonical Glx-Phe framework</u>

To evaluate whether the canonical Glx–Phe isotope framework correctly identified herbivores with atypical isotopic compositions, a linear discriminant analysis (LDA) was performed using

$\delta^{15}N$ values of Glx and Phe. Individuals previously identified as potential misclassifications in the hierarchical clustering analysis (including capybaras (*Hydrochoerus hydrochaeris*), agoutis (*Dasyprocta variegata*), squirrels (*Sciurus spadiceus* and *Tamiasciurus hudsonicus*), spiny rats (*Proechimys* spp.), Hoffmann's two-toed sloths (*Choloepus hoffmanni*), and West Indian manatees (*Trichechus manatus*)) were excluded from model training and treated as unknown.

The LDA model was trained on the remaining specimens with known dietary assignments, using Glx and Phe $\delta^{15}N$ values as predictor variables. The excluded herbivores were then classified with the fitted model, and posterior probabilities were calculated for each dietary category (table S2). This leave-out approach tested whether specimens with atypical Glx–Phe isotope values would still be recognized as primary consumers when evaluated solely within the canonical two-amino-acid framework. Although occasional consumption of animal-derived material has been reported for some of these taxa (e.g., manatees (*2*) and *Choloepus* (*3*)), these items constitute only a minor component of their diets. Consequently, these individuals were treated as obligate herbivores for the purposes of evaluating classification performance. All fourteen were classified as omnivores when treated as unknowns (table S2).

Ordinary Least Squares (OLS)

Ordinary least squares (OLS) regression was used to quantify the relationship between ΔGlx–Phe and ΔGlx–Thr for herbivores and secondary consumers. Interaction terms were used to test whether slopes differed significantly between trophic groups. Herbivores and secondary consumers occupied nearly orthogonal trajectories (∠ 95°) in ΔGlx–Phe versus ΔGlx–Thr space (OLS slopes: −0.55 and +2.21; interaction $p < 0.0001$), reflecting fundamentally different scaling relationships between these two isotopic offsets. Among secondary consumers, ΔGlx–Thr covaried positively with ΔGlx–Phe, consistent with previous observations that Thr $\delta^{15}N$ decreases with trophic level and dietary protein content. In contrast, herbivores exhibited a weak negative relationship , and variation in ΔGlx–Thr (~16.8‰) was nearly twice that observed for ΔGlx–Phe (~8.9‰), indicating that Thr captures an additional source of isotopic variation largely independent of the canonical trophic axis.

Standardized Major Axis (SMA) Regression

Relationships between ΔGlx–Phe and ΔGlx–Thr were evaluated using standardized major axis (SMA) regression implemented in the **smatr** R package. SMA was chosen because both isotopic variables contain measurement error and the objective was to characterize their scaling relationship rather than predict one variable from the other. Differences in slopes between groups were assessed using likelihood ratio tests implemented in **smatr**.

Three complementary analyses were performed. First, herbivores and secondary consumers were compared to determine whether the relationship between ΔGlx–Phe and ΔGlx–Thr differed among major trophic groups. Second, the analysis was repeated after excluding marine and freshwater piscivorous taxa to evaluate whether the observed relationships were driven by aquatic food webs. Third, herbivores were subdivided into foregut and hindgut fermenters to examine whether digestive physiology influenced the relationship between the two isotopic variables. As an additional sensitivity analysis, herbivore SMA regressions were repeated after excluding capybaras (*Hydrochoerus hydrochaeris*), manatees (*Trichechus manatus*), and koalas (*Phascolarctos cinereus*), which exhibited atypical Thr $\delta^{15}N$ values relative to other herbivores.

Herbivores and secondary consumers exhibited significantly different SMA slopes (likelihood ratio = 16.49, $P < 0.001$; Table S5). Herbivores showed a weak negative relationship between ΔGlx–Phe and ΔGlx–Thr (β = −1.45, 95% CI = −1.84 to −1.13, $R^2 = 0.15$), whereas secondary consumers exhibited a strong positive relationship (β = 2.58, 95% CI = 2.26–2.93, $R^2 = 0.74$).

Excluding marine and freshwater piscivorous taxa produced qualitatively identical results, Herbivores and secondary consumers continued to exhibit significantly different slopes (likelihood ratio = 9.81, $P = 0.002$), indicating that the observed scaling relationships are not driven by aquatic food webs.

Within herbivores, SMA slopes did not differ significantly between hindgut and foregut fermenters (likelihood ratio = 3.16, $P = 0.075$). Hindgut fermenters exhibited a negative relationship between ΔGlx–Phe and ΔGlx–Thr, whereas no significant relationship was detected among foregut fermenters. Excluding capybaras, manatees, and koalas did not qualitatively alter these results. A complete summary of all sensitivity analyses is summarized in Table S5.

Phylogenetic thinning

To evaluate whether the contrasting isotopic scaling relationships could arise from uneven sampling of closely related taxa, the SMA analysis was repeated after family-level phylogenetic thinning. Within herbivores and secondary consumers independently, one representative individual was randomly selected from each mammalian family. This procedure reduced the influence of repeatedly sampled families while preserving family representation within each trophic group and was repeated 1,000 times.

For each resampled dataset, standardized major axis (SMA) regressions were fit separately for herbivores and secondary consumers, and differences in slopes were evaluated using the common-slope likelihood ratio test implemented in smatr. The resulting distributions of slope estimates and common-slope $P$ values were used to assess the stability of the observed isotopic relationships under repeated family-level thinning.

Family-level thinning produced results highly consistent with those obtained from the complete dataset. Across all 1,000 resampled datasets, herbivore SMA slopes remained negative and secondary-consumer slopes remained positive in 100% of resampled datasets. Median slope estimates were −1.45 (95% resampling interval: −1.83 to −1.07) for herbivores and 2.59 (95% resampling interval: 2.45–2.75) for secondary consumers, nearly identical to the corresponding full-dataset estimates (−1.44 and 2.60, respectively). These results indicate that the contrasting isotopic scaling relationships are not driven by overrepresentation of particular mammalian families (Table S6).

Supplementary detail for Figure 3

Smooth curves connecting cluster medians were generated using shape-preserving piecewise cubic Hermite interpolation (PCHIP; scipy.interpolate.PchipInterpolator). The curves were used solely as visual guides and do not represent statistical model fits.

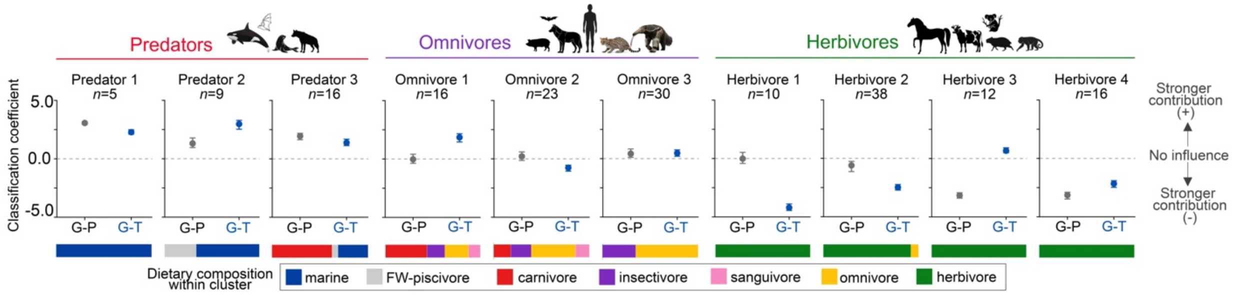

**TOP PREDATOR 1 (n=5)**
**Chiroptera:** *Myotis vivesi*

**TOP PREDATOR 2 (n=9)**
**Carnivora:** *Lontra canadensis, Pteronura brasiliensis, Arctocephalus pusillus*
**Cetacea:** *Delphinapterus leucas, Orcinus orca, Physeter macrocephalus, Leptonychotes weddellii,*

**TOP PREDATOR 3 (n= 16)**
**Carnivora:** *Crocuta crocuta, Arctocephalus pusillus*
**Cetacea:** *Balaena mysticetus, Balaenoptera physalus, Orcinus orca, Inia geoffrensis*
**Chiroptera:** *Trachops cirrhosus, Vampirum spectrum*
**Primates:** *Homo sapiens* (high marine protein consumption)

**OMNIVORE 1 (n= 16)**
**Carnivora:** *Galicitis vittata, Genetta genetta, Leopardus pardalis, Martes americana, Mephitis mephitis, Urocyon cinereoargentus*
**Chiroptera:** *Dipylla ecaudata, Desmodus rotundus*
**Didelphimorphia:** *Philander opossum*
**Macroscelidea:** *Elephantulus intufi*
**Pilosa:** *Myrmecophaga tridactyla*
**Primates:** *Saimiri brasiliensis*

**OMNIVORE 2 (n= 23)**
**Carnivora:** *Atelocynus* sp., *Galicitis vittata, Chrysocyon jubatus, Nasua nasua, Urocyon cinereoargentus*
**Chiroptera:** *Dipylla ecaudata, Desmodus rotundus*
**Cingulata:** *Dasypus novemcinctus, Chaetophractus vellerosus, Priodontes maximus*
**Didelphimorphia:** *Philander opossum*
**Eulipotyphla:** *Atelerix frontalis*
**Pilosa:** *Cyclopes didactylus*
**Primates:** *Saimiri brasiliensis, Homo neanderthalensis, Homo sapiens, Nycticebus coucang*
**Rodentia:** *Ondatra zibethicus*

**OMNIVORE 3 (n= 30)**
**Artiodactyla:** *Sus scrofa*
**Carnivora:** *Canis lupus, Potos flavus*
**Chiroptera:** *Glossophaga soricina, Lasionycteris noctivagans,Lasiurus cinereus*
**Cingulata:** *Euphractus sexcinctus*
**Pilosa:** *Tamandua sp., Cyclopes didactylus*
**Primates:** *Cebus apella, Saguinus fuscicollis, Homo sapiens (Hakemi site)*
**Rodentia:** *Glaucomys sabrinus, Rattus norvegicus, Myodes gapperi*

**HERBIVORE 1 (n= 10)**
**Rodentia:** *Dasyprocta variegata, Sciurus spadiceus, Dinomys branickii*
**Sirenia:** *Trichechus manaatus*

**HERBIVORE 2 (n= 38)**
**Artiodactyla:** *Bos* primigenius, *Syncerus caffer, Capra aegagrus, Pecari tajacu, Ovis aries*
**Pilosa:** *Bradypus variegatus, Choloepus hoffmanni*
**Rodentia:** *Dasyprocta variegata, Hydrochoeris hydrochaeris, Coendou bicolor, Proechimys brevicauda, P. quadruplicatus, Tamiasciurus hudsonicus*

**HERBIVORE 3 (n= 12)**
**Artiodactyla:** *Raphicerus campestris, Ovis aries*
**Diprotodontia:** *Phascolarctos cinereus*
**Primates:** *Aotus* spp.
**Rodentia:** *Coendou* sp.
**Pilosa:** *Bradypus variegatus*

**HERBIVORE 4 (n= 16)**
**Artiodactyla:** *Bos* spp., *Syncerus caffer, Giraffa giraffa, Capra aegagrus, Mazama chunyi, Pecari tajacu, Sus scrofa, Rangifer tarandus, Ovis aries*
**Chiroptera:** *Artibeus jamaicensis, Glossophaga soricina*
**Lagomorpha:** *Sylvilagus brasiliensis*
**Perissodactyla**: *Equus caballus*
**Primates:** *Ateles* sp., *Alouatta seniculus, Lagothrix lagotricha*
**Rodentia:** *Castor canadensis, Coendou* spp., *Myocastor coypus*

**Fig. S1. Standardized multinomial logistic regression coefficients for individual trophic clusters.** Hierarchical agglomerative clustering was repeated independently using ΔGlx–Phe and ΔGlx–Thr, and multinomial logistic regression models were fit using the resulting cluster assignments as response variables. Points (G–P= glutamic acid–phenylalanine $\delta^{15}$N offset, G–T= glutamic acid–threonine $\delta^{15}$N offset) show the distribution of standardized regression coefficients across 20 stratified bootstrap replicates for each predictor. Positive and negative coefficients indicate the direction of association with a given cluster, whereas larger absolute values indicate a greater contribution to discriminating that cluster from all others. This figure provides the complete taxonomic composition list of each cluster underlying the simplified version presented in Fig. 2.

A

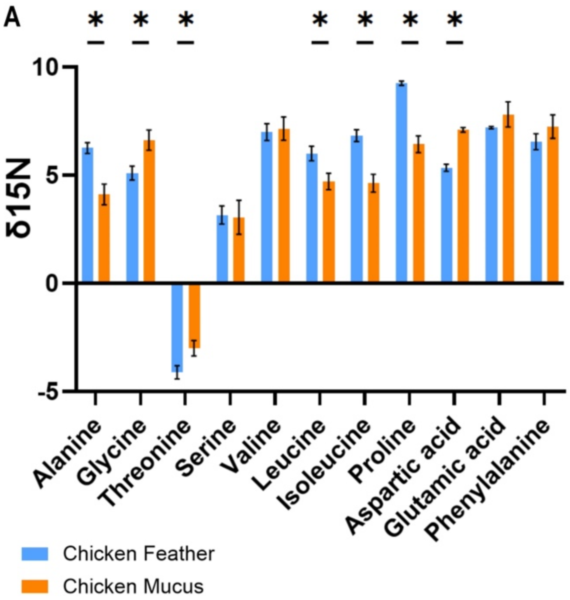

B

| | Chicken Feather | | Chicken Mucus | |
|---|---|---|---|---|
| | Mean | SD | Mean | SD |
| Alanine | 6.26 | 0.25 | 4.11 | 0.48 |
| Glycine | 5.10 | 0.32 | 6.62 | 0.47 |
| Threonine | -4.11 | 0.31 | -3.00 | 0.35 |
| Serine | 3.16 | 0.42 | 3.06 | 0.79 |
| Valine | 6.99 | 0.39 | 7.16 | 0.54 |
| Leucine | 6.01 | 0.34 | 4.71 | 0.38 |
| Isoleucine | 6.83 | 0.28 | 4.63 | 0.41 |
| Proline | 9.25 | 0.10 | 6.43 | 0.39 |
| Aspartic acid | 5.34 | 0.16 | 7.10 | 0.11 |
| Glutamic acid | 7.20 | 0.06 | 7.81 | 0.58 |
| Phenylalanine | 6.55 | 0.37 | 7.25 | 0.54 |

**Fig. S2. Pilot amino acid nitrogen isotope measurements of chicken intestinal mucus and feather. (A-B)** Measurements from one chicken individual show that Thr in intestinal mucus is $^{15}$N-enriched relative to feather, consistent with the mucin-retention hypothesis proposed here. Prior to mucus collection, the small intestinal lumen was gently flushed three times with 6 mL of phosphate-buffered saline (PBS). The intestine was then opened longitudinally and laid flat with the luminal surface facing upward. The luminal surface was scraped, and the content dried overnight at 60°C, mechanically homogenized, and stored at -20°C until derivatization and GC-IRMS analysis. Error bars represent the standard deviation (SD) of triplicate GC-IRMS measurements.

| Cluster | diet | Scientific_name_original |
|---|---|---|
| 0 | herb | H. hydrochaeris |
| 0 | herb | H. hydrochaeris |
| 0 | herb | H. hydrochaeris |
| 0 | herb | H. hydrochaeris |
| 0 | herb | Choloepus_hoffmanni |
| 0 | herb | Dasyprocta_variegata |
| 0 | herb | Dasyprocta_variegata |
| 0 | herb | Dasyprocta_variegata |
| 0 | herb | Dinomys_branickii |
| 0 | herb | Dinomys_branickii |
| 0 | mixed | Glaucomys_sabrinus |
| 0 | herb | Trichechus manatus |
| 0 | herb | Trichechus manatus |
| 0 | herb | Trichechus manatus |
| 0 | herb | P_brevicauda |
| 0 | herb | P_brevicauda |
| 0 | herb | P_brevicauda |
| 0 | mixed | Rattus_norvegicus |
| 0 | mixed | Rattus_norvegicus |
| 0 | herb | Tamiasciurus_hudsonicus |
| 0 | herb | Sciurus_spadiceus |
| 0 | herb | Sciurus_spadiceus |
| 0 | herb | Proechimys_quadruplicatus |
| 1 | mixed | Atelerix_frontalis |
| 1 | carn | Atelocynus_sp |
| 1 | carn | Atelocynus_sp |
| 1 | carn | Atelocynus_sp |
| 1 | sang | Diphylla ecaudata |
| 1 | sang | Diphylla ecaudata |
| 1 | sang | Desmodus rotundus |
| 1 | sang | Desmodus rotundus |
| 1 | mixed | Glossophaga soricina |
| 1 | mixed | Glossophaga soricina |
| 1 | insect | Lasiurus cinereus |
| 1 | insect | Lasiurus cinereus |
| 1 | insect | Lasiurus cinereus |
| 1 | insect | Lasionycteris noctivagans |
| 1 | insect | Lasionycteris noctivagans |
| 1 | insect | Lasionycteris noctivagans |
| 1 | mixed | Cebus_apella |
| 1 | mixed | Chaetophractus_vellerosus |
| 1 | carn | Chrysocyon_jubatus |
| 1 | insect | Cyclopes_didactylus |
| 1 | insect | Cyclopes_didactylus |
| 1 | mixed | Canis_lupus |
| 1 | mixed | Canis_lupus |
| 1 | insect | Euphractus_sexcinctus |
| 1 | carn | Galictis_vittata |
| 1 | carn | Genetta_genetta |
| 1 | mixed | Homo_neanderthalensis |
| 1 | mixed | Homo_neanderthalensis |
| 1 | mixed | Homo_neanderthalensis |
| 1 | mixed | Homo_sapiens_Hakemi |
| 1 | mixed | Homo_sapiens_RapaNui |
| 1 | mixed | Homo sapiens |
| 1 | mixed | Nycticebus_coucang |
| 1 | mixed | Sus_scrofa |
| 1 | insect | Priodontes_maximus |
| 1 | mixed | Rattus_norvegicus |
| 1 | mixed | Saimiri_boliviensis |
| 1 | insect | Tamandua_sp |
| 1 | insect | Tamandua_sp |
| 1 | mixed | Saguinus_fuscicollis |
| 1 | mixed | Saguinus_fuscicollis |
| 1 | mixed | Urocyon_cinereoargenteus |

| Cluster | diet | Scientific_name_original |
|---|---|---|
| 3 | herb | Aotus_sp |
| 3 | herb | Aotus_nigriceps |
| 3 | herb | Ateles_sp |
| 3 | herb | Bradypus_variegatus |
| 3 | herb | Syncerus_caffer |
| 3 | herb | Castor_canadensis |
| 3 | mixed | Cebus_apella |
| 3 | herb | Coendou_bicolor |
| 3 | herb | Coendou_bicolor |
| 3 | herb | Alouatta_seniculus |
| 3 | herb | Alouatta_seniculus |
| 3 | herb | Alouatta_seniculus |
| 3 | herb | Mazama_chunyi |
| 3 | herb | Myocastor_coypus |
| 3 | mixed | Myodes_gapperi |
| 3 | herb | Ovis_aries |
| 3 | herb | Raphicerus_campestris |
| 3 | herb | Raphicerus_campestris |
| 4 | marine | Delphinapterus_leucas |
| 4 | marine | Orcinus_orca |
| 4 | marine | Orcinus orca |
| 4 | fish | Lontra_canadensis |
| 4 | fish | Pteronura_brasiliensis |
| 4 | fish | Pteronura_brasiliensis |
| 4 | marine | Arctocephalus_pusillus |
| 4 | marine | Physeter_macrocephalus |
| 4 | marine | Physeter macrocephalus |
| 4 | marine | Leptonychotes weddellii |
| 5 | carn | Trachops cirrhosus |
| 5 | carn | Trachops cirrhosus |
| 5 | carn | Trachops cirrhosus |
| 5 | carn | Vampyrum spectrum |
| 5 | carn | Vampyrum spectrum |
| 5 | carn | Vampyrum spectrum |
| 5 | marine | Balaena_mysticetus |
| 5 | marine | Balaenoptera physalus |
| 5 | marine | Homo_sapiensHMP |
| 5 | carn | Crocuta_crocuta |
| 5 | carn | Crocuta_crocuta |
| 5 | carn | Crocuta_crocuta |
| 5 | carn | Crocuta_crocuta |
| 5 | marine | Orcinus_orca |
| 5 | marine | Arctocephalus_pusillus |
| 5 | marine | indet |
| 6 | marine | Myotis vivesi |
| 6 | marine | Myotis vivesi |
| 6 | marine | Myotis vivesi |
| 6 | marine | Myotis vivesi |
| 6 | marine | Myotis vivesi |

| Cluster | diet | Scientific_name_original |
|---|---|---|
| 7 | herb | Ateles_sp |
| 7 | herb | Artibeus jamaicensis |
| 7 | herb | Artibeus jamaicensis |
| 7 | herb | Artibeus jamaicensis |
| 7 | mixed | Glossophaga soricina |
| 7 | herb | Bos_taurus |
| 7 | herb | Bos_primigenius |
| 7 | herb | Bos_primigenius |
| 7 | herb | Bos sp |
| 7 | herb | Bos sp |
| 7 | herb | Bradypus_variegatus |
| 7 | herb | Bradypus_variegatus |
| 7 | herb | Bradypus_variegatus |
| 7 | herb | Syncerus_caffer |
| 7 | herb | Choloepus_hoffmanni |
| 7 | herb | Choloepus_hoffmanni |
| 7 | herb | Choloepus_hoffmanni |
| 7 | herb | Choloepus_hoffmanni |
| 7 | herb | Coendou |
| 7 | herb | Coendou_bicolor |
| 7 | herb | Giraffa_giraffa |
| 7 | herb | Capra_aegagrus |
| 7 | herb | Capra_aegagrus |
| 7 | herb | Capra_aegagrus |
| 7 | herb | Equus_caballus |
| 7 | herb | Equus_caballus |
| 7 | herb | Equus_caballus |
| 7 | mixed | Pecari_tajacu |
| 7 | mixed | Pecari_tajacu |
| 7 | mixed | Sus_scrofa |
| 7 | mixed | Sus_scrofa |
| 7 | mixed | Potos_flavus |
| 7 | mixed | Potos_flavus |
| 7 | herb | Rangifer_tarandus |
| 7 | herb | Ovis_aries |
| 7 | herb | Ovis_aries |
| 7 | herb | Ovis_aries |
| 7 | herb | Ovis_aries |
| 7 | herb | Sylvilagus_brasiliensis |
| 7 | herb | Sylvilagus_brasiliensis |
| 7 | herb | Lagothrix lagotricha |
| 7 | herb | Lagothrix lagotricha |
| 2 | mixed | Dasypus_novemcinctus |
| 2 | sang | Diphylla ecaudata |
| 2 | sang | Desmodus rotundus |
| 2 | insect | Elephantulus_intufi |
| 2 | carn | Galictis_vittata |
| 2 | mixed | Homo_sapiensC3 |
| 2 | mixed | Homo_sapiensC4 |
| 2 | fish | Inia_geoffrensis |
| 2 | carn | Leopardus_pardalis |
| 2 | carn | Leopardus_pardalis |
| 2 | carn | Leopardus_pardalis |
| 2 | mixed | Martes_americana |
| 2 | mixed | Martes_americana |
| 2 | mixed | Ondatra_zibethicus |
| 2 | insect | Myrmecophaga_tridactyla |
| 2 | insect | Myrmecophaga_tridactyla |
| 2 | mixed | Nasua_nasua |
| 2 | carn | Philander_opposum |
| 2 | insect | Philander |
| 2 | insect | Philander |
| 2 | mixed | Saimiri |
| 2 | mixed | Saimiri_brasiliensis |
| 2 | mixed | Mephitis_mephitis |
| 2 | carn | Urocyon_cinereoargentus |

**Table S1.**

Clustering based on Glx and Thr normalized to Phe (threshold=20). Highlighted, notorious inconsistencies with results obtained with Thr and Phe normalized to Glx.

| Taxon | Phe | Glx | Glx-Phe | Predicted_diet | Prob_carn | Prob_fish | Prob_herb | Prob_insec | Prob_mari | Prob_omni | Prob_sang | Max_probability |
|---|---|---|---|---|---|---|---|---|---|---|---|---|
| H. hydrochaeris | 9.894 | 13.66 | 3.766 | omnivore | 0.02 | 0 | 0.105 | 0.237 | 0 | 0.596 | 0.042 | 0.596 |
| H. hydrochaeris | 6.304 | 9.873 | 3.569 | omnivore | 0.029 | 0 | 0.133 | 0.157 | 0 | 0.664 | 0.017 | 0.664 |
| H. hydrochaeris | 7.16 | 10.581 | 3.421 | omnivore | 0.022 | 0 | 0.148 | 0.166 | 0 | 0.644 | 0.019 | 0.644 |
| C. hoffmanni | 8.075 | 11.721 | 3.646 | omnivore | 0.024 | 0 | 0.122 | 0.193 | 0 | 0.635 | 0.026 | 0.635 |
| D. variegata | 14.082 | 17.273 | 3.191 | omnivore | 0.005 | 0.001 | 0.147 | 0.299 | 0 | 0.479 | 0.069 | 0.479 |
| Proechimys | 8.384 | 11.946 | 3.562 | omnivore | 0.021 | 0 | 0.13 | 0.196 | 0 | 0.627 | 0.027 | 0.627 |
| Proechimys | 7.916 | 10.642 | 2.726 | omnivore | 0.009 | 0 | 0.247 | 0.146 | 0 | 0.584 | 0.014 | 0.584 |
| Proechimys | 8.044 | 10.943 | 2.899 | omnivore | 0.011 | 0 | 0.218 | 0.157 | 0 | 0.597 | 0.016 | 0.597 |
| Tamiasciurus | 4.535 | 9.042 | 4.507 | omnivore | 0.092 | 0 | 0.059 | 0.151 | 0 | 0.678 | 0.019 | 0.678 |
| Proechimys | 6.728 | 9.619 | 2.891 | omnivore | 0.013 | 0 | 0.223 | 0.136 | 0 | 0.616 | 0.012 | 0.616 |
| Dasyprocta | 5.514 | 8.848 | 3.334 | omnivore | 0.026 | 0 | 0.162 | 0.135 | 0 | 0.665 | 0.012 | 0.665 |
| Trichechus mantus | 8.232 | 11.132 | 2.9 | omnivore | 0.011 | 0 | 0.218 | 0.16 | 0 | 0.594 | 0.017 | 0.594 |
| Trichechus mantus | 9.533 | 12.423 | 2.89 | omnivore | 0.009 | 0 | 0.214 | 0.183 | 0 | 0.572 | 0.022 | 0.572 |
| S. spadiceus | 7.842 | 11.14 | 3.298 | omnivore | 0.018 | 0 | 0.162 | 0.173 | 0 | 0.627 | 0.02 | 0.627 |

**Table S2.**

Linear discriminant analysis trained on Glx–Phe $\delta^{15}N$ (excluding these individuals from the training dataset) classified fourteen herbivores as omnivores when treated as unknown.

| Cluster | diet | Scientific_name_original | Cluster | diet | Scientific_name_original | Cluster | diet | Scientific_name_original |
|---|---|---|---|---|---|---|---|---|
| 1 | mixed | Glossophaga soricina | 3 | herb | Aotus_sp | 7 | carn | Atelocynus_sp |
| 1 | mixed | Glossophaga soricina | 3 | herb | Aotus_nigriceps | 7 | sang | Diphylla ecaudata |
| 1 | insect | Lasiurus cinereus | 3 | herb | Ateles_sp | 7 | sang | Desmodus rotundus |
| 1 | insect | Lasiurus cinereus | 3 | herb | Artibeus jamaicensis | 7 | sang | Desmodus rotundus |
| 1 | insect | Lasiurus cinereus | 3 | herb | Artibeus jamaicensis | 7 | carn | Chrysocyon_jubatus |
| 1 | insect | Lasionycteris noctivagans | 3 | herb | Artibeus jamaicensis | 7 | carn | Galictis_vittata |
| 1 | insect | Lasionycteris noctivagans | 3 | mixed | Glossophaga soricina | 7 | mixed | Homo sapiens |
| 1 | insect | Lasionycteris noctivagans | 3 | herb | Bos_taurus | 7 | fish | Inia_geoffrensis |
| 1 | herb | H. hydrochaeris | 3 | herb | Bos_primigenius | 7 | carn | Leopardus_pardalis |
| 1 | herb | H. hydrochaeris | 3 | herb | Bos sp | 7 | carn | Leopardus_pardalis |
| 1 | herb | H. hydrochaeris | 3 | herb | Bos sp | 7 | carn | Leopardus_pardalis |
| 1 | herb | H. hydrochaeris | 3 | herb | Bradypus_variegatus | 7 | mixed | Martes_americana |
| 1 | mixed | Cebus_apella | 3 | herb | Bradypus_variegatus | 7 | mixed | Nasua_nasua |
| 1 | herb | Choloepus_hoffmanni | 3 | herb | Coendou | 7 | carn | Urocyon_cinereoargentus |
| 1 | herb | Choloepus_hoffmanni | 3 | herb | Coendou_bicolor | 8 | herb | Ateles_sp |
| 1 | herb | Dasyprocta_variegata | 3 | herb | Coendou_bicolor | 8 | herb | Syncerus_caffer |
| 1 | herb | Dasyprocta_variegata | 3 | herb | Dinomys_branickii | 8 | herb | Castor_canadensis |
| 1 | herb | Dasyprocta_variegata | 3 | herb | Capra_aegagrus | 8 | herb | Alouatta_seniculus |
| 1 | mixed | Canis_lupus | 3 | herb | Equus_caballus | 8 | herb | Alouatta_seniculus |
| 1 | mixed | Canis_lupus | 3 | herb | Equus_caballus | 8 | herb | Alouatta_seniculus |
| 1 | mixed | Homo_sapiens_Hakemi | 3 | herb | Equus_caballus | 8 | herb | Mazama_chunyi |
| 1 | herb | Trichechus manatus | 3 | mixed | Sus_scrofa | 8 | herb | Myocastor_coypus |
| 1 | herb | Trichechus manatus | 3 | herb | Rangifer_tarandus | 9 | marine | Delphinapterus_leucas |
| 1 | mixed | Martes_americana | 3 | herb | Ovis_aries | 9 | marine | Orcinus_orca |
| 1 | mixed | Ondatra_zibethicus | 3 | herb | Ovis_aries | 0 | carn | Vampyrum spectrum |
| 1 | insect | Myrmecophaga_tridactyla | 3 | herb | Ovis_aries | 0 | marine | Balaena_mysticetus |
| 1 | mixed | Sus_scrofa | 3 | herb | Ovis_aries | 0 | marine | Homo_sapiensHMP |
| 1 | mixed | Potos_flavus | 3 | herb | Raphicerus_campestris | 0 | carn | Crocuta_crocuta |
| 1 | indet | P_brevicauda | 3 | herb | Sylvilagus_brasiliensis | 0 | carn | Crocuta_crocuta |
| 1 | indet | P_brevicauda | 3 | herb | Sylvilagus_brasiliensis | 0 | marine | Orcinus orca |
| 1 | indet | P_brevicauda | 3 | herb | Lagothrix lagotricha | 0 | fish | Lontra_canadensis |
| 1 | mixed | Rattus_norvegicus | 3 | herb | Lagothrix lagotricha | 0 | marine | Orcinus_orca |
| 1 | mixed | Saimiri | 4 | marine | Myotis vivesi | 0 | marine | Arctocephalus_pusillus |
| 1 | herb | Sciurus_spadiceus | 4 | marine | Myotis vivesi | 0 | marine | Physeter_macrocephalus |
| 1 | herb | Sciurus_spadiceus | 4 | marine | Myotis vivesi | 0 | marine | Physeter macrocephalus |
| 1 | herb | Proechimys_quadruplicatus | 4 | marine | Myotis vivesi | 0 | marine | Leptonychotes weddellii |
| 1 | mixed | Saguinus_fuscicollis | 4 | marine | Myotis vivesi | 0 | marine | whale |
| 1 | mixed | Saguinus_fuscicollis | 5 | herb | Bos_primigenius | | | |
| 2 | mixed | Dasypus_novemcinctus | 5 | herb | Bradypus_variegatus | | | |
| 2 | mixed | Atelerix_frontalis | 5 | herb | Bradypus_variegatus | | | |
| 2 | carn | Atelocynus_sp | 5 | herb | Syncerus_caffer | | | |
| 2 | carn | Atelocynus_sp | 5 | mixed | Cebus_apella | | | |
| 2 | sang | Diphylla ecaudata | 5 | herb | Choloepus_hoffmanni | | | |
| 2 | sang | Diphylla ecaudata | 5 | herb | Choloepus_hoffmanni | | | |
| 2 | sang | Desmodus rotundus | 5 | herb | Choloepus_hoffmanni | | | |
| 2 | mixed | Chaetophractus_vellerosus | 5 | herb | Coendou_bicolor | | | |
| 2 | insect | Cyclopes_didactylus | 5 | herb | Dinomys_branickii | | | |
| 2 | insect | Cyclopes_didactylus | 5 | herb | Giraffa_giraffa | | | |
| 2 | insect | Elephantulus_intufi | 5 | herb | Capra_aegagrus | | | |
| 2 | insect | Euphractus_sexcinctus | 5 | herb | Capra_aegagrus | | | |
| 2 | mixed | Glaucomys_sabrinus | 5 | mixed | Homo_sapiensC4 | | | |
| 2 | mixed | Homo_neanderthalensis | 5 | herb | Trichechus manatus | | | |
| 2 | mixed | Homo_neanderthalensis | 5 | mixed | Pecari_tajacu | | | |
| 2 | mixed | Homo_neanderthalensis | 5 | mixed | Pecari_tajacu | | | |
| 2 | mixed | Homo_sapiensC3 | 5 | mixed | Sus_scrofa | | | |
| 2 | mixed | Homo_sapiens_RapaNui | 5 | mixed | Potos_flavus | | | |
| 2 | mixed | Nycticebus_coucang | 5 | mixed | Myodes_gapperi | | | |
| 2 | insect | Myrmecophaga_tridactyla | 5 | herb | Ovis_aries | | | |
| 2 | carn | Philander_opposum | 5 | herb | Raphicerus_campestris | | | |
| 2 | insect | Philander | 6 | carn | Trachops cirrhosus | | | |
| 2 | insect | Philander | 6 | carn | Trachops cirrhosus | | | |
| 2 | insect | Priodontes_maximus | 6 | carn | Trachops cirrhosus | | | |
| 2 | mixed | Rattus_norvegicus | 6 | carn | Vampyrum spectrum | | | |
| 2 | mixed | Rattus_norvegicus | 6 | carn | Vampyrum spectrum | | | |
| 2 | herb | Tamiasciurus_hudsonicus | 6 | marine | Balaenoptera physalus | | | |
| 2 | mixed | Saimiri_boliviensis | 6 | carn | Galictis_vittata | | | |
| 2 | mixed | Saimiri_brasiliensis | 6 | carn | Genetta_genetta | | | |
| 2 | mixed | Mephitis_mephitis | 6 | carn | Crocuta_crocuta | | | |
| 2 | insect | Tamandua_sp | 6 | carn | Crocuta_crocuta | | | |
| 2 | insect | Tamandua_sp | 6 | fish | Pteronura_brasiliensis | | | |
| 2 | mixed | Urocyon_cinereoargenteus | 6 | fish | Pteronura_brasiliensis | | | |
| | | | 6 | marine | Arctocephalus_pusillus | | | |

**Table S3.**

Unsupervised clustering results based on canonical Glx–Phe $\delta^{15}N$ framework. Systematic misclassification of herbivores and secondary consumers highlighted in pink.

**HERBIVORES**

| Taxon | Scientific_name_( | diet | Glx | Phe | Thr | D_Glx_Phe | D_Glx_Thr |
|---|---|---|---|---|---|---|---|
| Dasyprocta1 | Dasyprocta_varieg | herb | 8.84838678 | 5.51368878 | 0.74272944 | 3.334698 | 8.10565734 |
| Dasyprocta3 | Dasyprocta_varieg | herb | 10.0510358 | 7.96992538 | 1.56934491 | 2.08111042 | 8.48169089 |
| Dinomys1 | Dinomys_branicki | herb | 4.92417074 | 6.00642218 | -3.1926527 | -1.0822514 | 8.11682347 |
| Dinomys2 | Dinomys_branicki | herb | 3.96136363 | 4.16693584 | -6.5431788 | -0.2055722 | 10.5045424 |
| manatee1 | Trichechus manat | herb | 11.1320274 | 8.23238116 | 3.11505183 | 2.89964621 | 8.01697554 |
| manatee2 | Trichechus manat | herb | 17.3233623 | 16.1719549 | 7.64177468 | 1.15140744 | 9.68158764 |
| manatee3 | Trichechus manat | herb | 12.4229892 | 9.5325958 | 4.49026087 | 2.89039336 | 7.93272828 |
| Sciurus1 | Sciurus_spadiceus | herb | 11.1395545 | 7.84185759 | 8.00486476 | 3.29769695 | 3.13468978 |
| Sciurus2 | Sciurus_spadiceus | herb | 10.0194205 | 7.6135599 | 5.86283997 | 2.40586055 | 4.15658048 |
| capibara1 | H. hydrochaeris | herb | 11.5134226 | 9.06691558 | -1.4467394 | 2.44650701 | 12.960162 |
| capibara2 | H. hydrochaeris | herb | 13.659746 | 9.89413737 | 1.0288421 | 3.76560865 | 12.6309039 |
| capibara3 | H. hydrochaeris | herb | 9.87322494 | 6.30355346 | -1.1339632 | 3.56967148 | 11.0071881 |
| capibara4 | H. hydrochaeris | herb | 10.581458 | 7.16016696 | -1.8914083 | 3.42129099 | 12.4728663 |
| Choloepus2 | Choloepus_hoffma | herb | 11.7210401 | 8.07512166 | -3.1470106 | 3.64591842 | 14.8680507 |
| Dasyprocta2 | Dasyprocta_varieg | herb | 17.272744 | 14.0821146 | 2.19655743 | 3.19062933 | 15.0761866 |
| Proechimys1 | P_brevicauda | herb | 11.9462747 | 8.38438607 | -2.6818964 | 3.56188867 | 14.6281711 |
| Proechimys2 | P_brevicauda | herb | 10.6419933 | 7.91588046 | -4.1457419 | 2.72611281 | 14.7877352 |
| Proechimys3 | P_brevicauda | herb | 10.9428827 | 8.04447092 | -2.4207183 | 2.89841181 | 13.3636011 |
| red squirrel | Tamiasciurus_hud | herb | 9.042026 | 4.53497514 | -4.0150863 | 4.50705085 | 13.0571123 |
| spiny rat | Proechimys_quad | herb | 9.6188023 | 6.72815296 | -4.2988433 | 2.89064934 | 13.9176456 |
| Aotus | Aotus_sp | herb | 9.82013072 | 11.3509295 | -10.10381 | -1.5307988 | 19.923941 |
| Aotus_nigric | Aotus_nigriceps | herb | 14.7355975 | 16.099182 | -4.7293984 | -1.3635845 | 19.4649959 |
| Bradypus | Bradypus_variega | herb | 10.7660317 | 12.6600043 | -6.5718786 | -1.8939726 | 17.3379103 |
| Coendou2 | Coendou_bicolor | herb | 8.70504542 | 10.1441176 | -7.5962785 | -1.4390721 | 16.3013239 |
| koala1 | Phascolarctos cine | herb | 8.17063593 | 10.8453457 | -11.332292 | -2.6747098 | 19.502928 |
| koala2 | Phascolarctos cine | herb | 6.29203124 | 9.13726331 | -12.716413 | -2.8452321 | 19.0084442 |
| koala3 | Phascolarctos cine | herb | 8.43801317 | 11.3769871 | -10.826998 | -2.9389739 | 19.2650109 |
| koala4 | Phascolarctos cine | herb | 7.81703459 | 11.3559798 | -11.252682 | -3.5389452 | 19.0697165 |
| koala5 | Phascolarctos cine | herb | 9.37790673 | 12.9230213 | -8.8351667 | -3.5451146 | 18.2130735 |
| Sheep5 | Ovis_aries | herb | 15.7 | 16.2 | -2.4 | -0.5 | 18.1 |
| steenbok1 | Raphicerus_camp | herb | 14.4 | 15 | -4 | -0.6 | 18.4 |
| steenbok2 | Raphicerus_camp | herb | 13.8 | 14.1 | -3.5 | -0.3 | 17.3 |
| Ateles | Ateles_sp | herb | 7.43897762 | 9.26808296 | -6.8064497 | -1.8291053 | 14.2454273 |
| Ateles2 | Ateles_sp | herb | 6.9508181 | 9.7962925 | -3.0135817 | -2.8454744 | 9.96439981 |
| Bos | Bos_taurus | herb | 12 | 12.6 | -2.4 | -0.6 | 14.4 |
| Bradypus1 | Bradypus_variega | herb | 14.54 | 14.1 | -1.03 | 0.44 | 15.57 |
| Bradypus2 | Bradypus_variega | herb | 14.61 | 15.34 | 0.28 | -0.73 | 14.33 |
| Bradypus3 | Bradypus_variega | herb | 14.0796676 | 12.9478421 | 0.73649726 | 1.13182558 | 13.3431704 |
| buffalo1 | Syncerus_caffer | herb | 8.5 | 8.7 | -6.6 | -0.2 | 15.1 |
| buffalo2 | Syncerus_caffer | herb | 3.3 | 7.7 | -10 | -4.4 | 13.3 |
| castor | Castor_canadensi | herb | 6.72771094 | 9.47590287 | -6.8850178 | -2.7481919 | 13.6127288 |
| Choloepus | Choloepus_hoffma | herb | 11.413257 | 10.8237694 | -5.1230303 | 0.5894876 | 16.5362873 |
| Choloepus3 | Choloepus_hoffma | herb | 14.1091554 | 13.9866598 | -0.0983117 | 0.12249562 | 14.2074671 |
| Choloepus4 | Choloepus_hoffma | herb | 10.5307248 | 8.46244175 | -5.4647659 | 2.06828301 | 15.9954906 |
| Choloepus5 | Choloepus_hoffma | herb | 8.9080219 | 8.39281057 | -5.7608356 | 0.51521133 | 14.6688575 |
| Coendou1 | Coendou | herb | 6.67937747 | 8.12408687 | -5.6141396 | -1.4447094 | 12.2935171 |
| Coendou3 | Coendou_bicolor | herb | 9.35525055 | 11.1446558 | -5.5401481 | -1.7894052 | 14.8953987 |
| Coendou4 | Coendou_bicolor | herb | 10.0864748 | 8.63482017 | -4.4573782 | 1.45165459 | 14.5438529 |
| giraffa | Giraffa_giraffa | herb | 9.99927818 | 9.96828216 | -2.6600119 | 0.03099602 | 12.6592901 |
| howler_mon | Alouatta_seniculu | herb | 8.9515249 | 12.9343302 | -1.7471388 | -3.9828053 | 10.6986637 |
| howler_mon | Alouatta_seniculu | herb | 9.17309439 | 13.5777868 | -1.865047 | -4.4046924 | 11.0381414 |
| howler_mon | Alouatta_seniculu | herb | 9.40147813 | 13.3670124 | -5.16052 | -3.9655343 | 14.5619981 |
| LA_squirrel | Sciurus_niger | herb | 9.23507193 | 7.0085105 | -5.2204255 | 2.22656143 | 14.4554975 |
| Mazama | Mazama_chunyi | herb | 5.05876116 | 8.21137655 | -9.7018154 | -3.1526154 | 14.7605766 |
| myocastor | Myocastor_coypus | herb | 9.29887414 | 11.8137419 | -2.9158851 | -2.5148677 | 12.2147592 |
| pecari1 | Pecari_tajacu | mixed | 7.84231182 | 6.86047319 | -4.8263136 | 0.98183863 | 12.6686255 |
| pecari2 | Pecari_tajacu | mixed | 10.1825472 | 9.15899834 | -5.8144372 | 1.02354891 | 15.9969845 |
| Sheep4 | Ovis_aries | herb | 14.7 | 13.4 | -0.4 | 1.3 | 15.1 |
| Sylvilagus | Sylvilagus_brasilie | herb | 8.30817546 | 8.87830682 | -5.2594289 | -0.5701314 | 13.5676043 |
| Sylvilagus2 | Sylvilagus_brasilie | herb | 7.02445812 | 7.49494382 | -6.739952 | -0.4704857 | 13.7644101 |
| woolly_monk | Lagothrix lagotricl | herb | 6.43408871 | 7.62629722 | -5.4981273 | -1.1922085 | 11.9322161 |
| woolly_monk | Lagothrix lagotricl | herb | 7.96427216 | 8.83713866 | -5.7969377 | -0.8728665 | 13.7612099 |

**OMNIVORES**

| Taxon | Scientific_na | diet | Glx | Phe | Thr | D_Glx_Phe | D_Glx_Thr |
|---|---|---|---|---|---|---|---|
| elephant-shr | Elephantulus | insect | 18.5742279 | 13.0994962 | -11.734701 | 5.47473169 | 30.3089285 |
| Galictis2 | Galictis_vitta | carn | 16.0404165 | 6.31997964 | -13.657695 | 9.72043686 | 29.6981114 |
| Genetta | Genetta_ger | carn | 20.87 | 10.19 | -10.05 | 10.68 | 30.92 |
| Leopardus1 | Leopardus_p | carn | 15.3127174 | 8.72943789 | -17.45428 | 6.58327952 | 32.7669979 |
| Leopardus2 | Leopardus_p | carn | 15.848682 | 8.16444961 | -15.457786 | 7.68423238 | 31.3064684 |
| Leopardus3 | Leopardus_p | carn | 14.2965259 | 7.48809087 | -16.82813 | 6.80843504 | 31.1246559 |
| Martes | Martes_ame | mixed | 11.05 | 8.67 | -19.49 | 2.38 | 30.54 |
| Martes | Martes_ame | mixed | 14.1550673 | 6.85590972 | -15.875553 | 7.29915755 | 30.0306198 |
| Myrmecopha | Myrmecopha | insect | 15.7623467 | 11.6050936 | -17.570799 | 4.15725317 | 33.3331455 |
| Myrmecopha | Myrmecopha | insect | 13.6984687 | 8.84826234 | -19.222098 | 4.85020634 | 32.920567 |
| opossum | Philander_op | carn | 13.7363015 | 7.67474327 | -17.425678 | 6.06155821 | 31.1619797 |
| Saimiri2 | Saimiri_bras | mixed | 14.1324599 | 9.58544979 | -19.470487 | 4.54701008 | 33.6029468 |
| skunk | Mephitis_me | mixed | 11.6905045 | 6.92782203 | -19.868825 | 4.76268244 | 31.5593299 |
| Urocyon2 | Urocyon_cine | carn | 14.0876552 | 7.08597505 | -16.728288 | 7.00168017 | 30.8159427 |
| 9b-armadillo | Dasypus_nov | mixed | 17.613157 | 12.3021284 | -10.961793 | 5.31102862 | 28.5749498 |
| Atelerix | Atelerix_fron | mixed | 13.53 | 7.4 | -12.98 | 6.13 | 26.51 |
| Atelocynus2 | Atelocynus_s | carn | 16.6381565 | 11.2968957 | -10.018645 | 5.34126078 | 26.6568013 |
| Atelocynus3 | Atelocynus_s | carn | 15.338922 | 8.05184428 | -10.726704 | 7.28707771 | 26.0656258 |
| Chrysocyon1 | Chrysocyon_j | carn | 13.54 | 6.32 | -13.07 | 7.22 | 26.61 |
| Galictis1 | Galictis_vitta | carn | 18.5715313 | 11.8131903 | -10.505314 | 6.75834101 | 29.0768454 |
| HomoC3 | Homo_sapie | mixed | 12.3 | 7.7 | -15.1 | 4.6 | 27.4 |
| HomoC4 | Homo_sapie | mixed | 11.1 | 9.9 | -14.6 | 1.2 | 25.7 |
| Hsapiens (on | Homo sapien | mixed | 14.4526344 | 7.27645083 | -13.72461 | 7.17618353 | 28.1772446 |
| muskrat | Ondatra_zibe | mixed | 12.3127195 | 9.03671912 | -14.456267 | 3.27600041 | 26.7689867 |
| Nasua | Nasua_nasua | mixed | 16.09 | 9.11 | -13.02 | 6.98 | 29.11 |
| Philander1 | Philander | insect | 12.0360289 | 7.21692481 | -15.271709 | 4.81910407 | 27.3077375 |
| Philander2 | Philander | insect | 15.2828394 | 10.6983448 | -12.23362 | 4.58449461 | 27.5164592 |
| Saimiri1 | Saimiri | mixed | 13.3755316 | 9.89878155 | -14.212402 | 3.47675008 | 27.5879335 |
| Atelocynus1 | Atelocynus_s | carn | 15.2483377 | 10.7148799 | -9.7911396 | 4.53345785 | 25.0394774 |
| Chaetophrac | Chaetophrac | mixed | 10.86 | 4.63 | -13.26 | 6.23 | 24.12 |
| Cyclopes | Cyclopes_did | insect | 12.7744408 | 7.93124428 | -10.183099 | 4.84319648 | 22.9575396 |
| Cyclopes2 | Cyclopes_did | insect | 13.3523262 | 7.16592024 | -10.928776 | 6.18640601 | 24.2811019 |
| Euphractus | Euphractus_s | insect | 15.1182403 | 10.7454607 | -6.0885121 | 4.37277966 | 21.2067525 |
| HomoRapa | Homo_sapie | mixed | 17.76 | 11.69 | -6.26 | 6.07 | 24.02 |
| Hsapiens (ve | Homo sapien | vegan | 10.7652425 | 4.50115083 | -9.2054805 | 6.26409171 | 19.9707231 |
| Hsapiens (ve | Homo sapien | vegetarian | 13.4212798 | 7.78895317 | -8.7269006 | 5.63232664 | 22.1481804 |
| loris | Nycticebus_c | mixed | 14.7040998 | 10.3130444 | -8.9259568 | 4.39105539 | 23.6300566 |
| Priodontes | Priodontes_r | insect | 11.1664076 | 5.09698575 | -12.877441 | 6.06942188 | 24.0438486 |
| Rat3 | Rattus_norve | mixed | 15 | 10.89 | -4.75 | 4.11 | 19.75 |
| Saimiri | Saimiri_boliv | mixed | 12.74 | 6.69 | -11.18 | 6.05 | 23.92 |
| Tamandua1 | Tamandua_s | insect | 10.6891768 | 4.93842734 | -10.862639 | 5.75074949 | 21.5518162 |
| Tamandua2 | Tamandua_s | insect | 13.5816632 | 8.66541869 | -6.7241022 | 4.91624447 | 20.3057654 |
| tamarin2 | Saguinus_fus | mixed | 15.0004008 | 11.6118128 | -7.1508209 | 3.38858794 | 22.1512217 |
| Urocyon | Urocyon_cine | mixed | 14.12 | 8.54 | -10.68 | 5.58 | 24.8 |
| Cebus1 | Cebus_apella | mixed | 10.5268295 | 8.97362165 | -10.094935 | 1.55320781 | 20.6217643 |
| Cebus2 | Cebus_apella | mixed | 12.036914 | 9.6581053 | -6.8097024 | 2.37880873 | 18.8466164 |
| flying-squirre | Glaucomys_s | mixed | 16.2404374 | 10.4494118 | 0.11448482 | 5.79102559 | 16.1259526 |
| Potos1 | Potos_flavus | mixed | 10.9520924 | 7.85600372 | -6.4338112 | 3.09608867 | 17.3859035 |
| Potos2 | Potos_flavus | mixed | 10.7782801 | 9.06441489 | -7.2300166 | 1.71386517 | 18.0082967 |
| Rat1 | Rattus_norve | mixed | 12.17 | 7.19 | -5.81 | 4.98 | 17.98 |
| Rat2 | Rattus_norve | mixed | 17.54 | 12.99 | 0.19 | 4.55 | 17.35 |
| red vole | Myodes_gap | mixed | 10.9154316 | 10.2309146 | -8.1573392 | 0.68451703 | 19.0727708 |
| tamarin | Saguinus_fus | mixed | 14.8615587 | 12.7741703 | -4.1352628 | 2.08738845 | 18.9968215 |

**PREDATORS**

| Taxon | Scientific_na | diet | Glx | Phe | Thr | D_Glx_Phe | D_Glx_Thr |
|---|---|---|---|---|---|---|---|
| killer whale | Orcinus orca | marine | 26.19 | 12.28 | -17.1 | 13.91 | 43.29 |
| Lontra | Lontra_canad | fish | 24.914078 | 10.8805459 | -22.207505 | 14.0335321 | 47.1215832 |
| Pteronura1 | Pteronura_br | fish | 25.170861 | 13.6099793 | -24.348016 | 11.5608818 | 49.5188767 |
| Pteronura2 | Pteronura_br | fish | 21.272308 | 10.0803657 | -22.251287 | 11.1919423 | 43.5235954 |
| seal2 | Arctocephalu | marine | 25.4 | 10.7 | -27.1 | 14.7 | 52.5 |
| sperm-whale | Physeter mac | marine | 23.3333333 | 7.81666667 | -34.25 | 15.5166667 | 57.5833333 |
| weddell_sea | Leptonychote | marine | 21.5 | 5.9 | -30.65 | 15.6 | 52.15 |
| fin whale | Balaenoptera | marine | 15 | 4.45 | -22.15 | 10.55 | 37.15 |
| HomoHMP | Homo_sapie | marine | 19.4 | 7.3 | -20.6 | 12.1 | 40 |
| Inia | Inia_geoffrer | fish | 21.6724776 | 13.930434 | -18.41 | 7.74204363 | 40.0824776 |
| seal1 | Arctocephalu | marine | 22.6 | 11.2 | -18.1 | 11.4 | 40.7 |
| whaleRapaN | indet | marine | 14.24 | 0.71 | -23.31 | 13.53 | 37.55 |

**Table S4.** Unsupervised clustering results based on Glx-Phe and Glx-Thr $\delta^{15}N$ of measured data only (i.e., excluding imputed values). The clustering structure was largely consistent, especially among herbivores. The main observed difference was an increased in the number of predator clusters.

| Analysis | Group | n | SMA slope | 95% CI | Common slope LR | P |
|---|---|---|---|---|---|---|
| Main | Herbivore | 59 | -1.45 | -1.84 to -1.13 | 16.49 | <0.001 |
| | Secondary consumer | 65 | 2.58 | 2.26 to 2.93 | | |
| No marine | Herbivore | 59 | -1.45 | -1.84 to -1.13 | 9.81 | 0.002 |
| | Secondary consumer | 54 | 2.48 | 1.96 to 3.13 | | |
| Fermentation | Hindgut | 32 | -1.56 | -2.11 to -1.15 | 3.16 | 0.075 |
| | Foregut | 18 | 0.91 | 0.55 to 1.52 | | |
| Excluding taxa | Hindgut | 20 | -1.57 | -2.50 to -0.99 | 2.43 | 0.119 |
| | Foregut | 18 | 0.91 | 0.55 to 1.52 | | |

**Table S5. Sensitivity analyses of standardized major axis (SMA) regressions.** Standardized major axis (SMA) regression parameters for the relationship between ΔGlx–Thr and ΔGlx–Phe across the principal analyses presented in the study. Results are shown for the full dataset, after excluding marine and fish–consumer taxa (non-marine), among herbivores grouped by fermentation strategy (hindgut versus foregut), and after excluding herbivore taxa with atypical isotopic signatures (i.e., koalas, capybaras, and manatees). Reported values include sample size (*n*), SMA slope with 95% confidence intervals, and the likelihood-ratio (LR) test for differences in slopes between groups. The consistent sign and magnitude of SMA slopes across analyses demonstrate that the contrasting isotopic scaling relationships are robust to taxonomic composition and analytical assumptions.

| Metric | Herbivores | Secondary consumers |
|---|---|---|
| Full-dataset SMA slope | −1.44 | 2.60 |
| Median family-thinned slope | −1.45 | 2.59 |
| 95% resampling interval | −1.83 to −1.07 | 2.45 to 2.75 |

**Table S6. Robustness of SMA slopes to family-level phylogenetic thinning.**
Full-dataset slopes are compared with median slopes and 95% resampling intervals from 1000 family-thinned datasets.

**Data S1 (Excel file).** Compiled and newly generated amino acid $\delta^{15}N$ measurements, including specimen metadata and all variables used in the statistical analyses. The workbook contains two worksheets: **All_data**, which includes all 175 specimens and their associated metadata and amino acid isotope measurements; and **woNA**, which excludes specimens lacking threonine (Thr) $\delta^{15}N$ measurements, for sensitivity analyses requiring complete Thr data.

**Data S2 (TXT file).** Python scripts used to perform hierarchical clustering and multinomial logistic regression analyses of feature contributions.
